# Raw materials and manufacturing environment as determinants of miso microbial community

**DOI:** 10.1101/2024.10.09.614917

**Authors:** Kohei Ito, Marin Yamaguchi

## Abstract

Miso is a Japanese traditional fermented food with soybeans, salt and *koji*, and has gained attention among people for its sophisticated flavor and preservability. *Koshu miso* is a unique miso made by mixing two types of *koji* (rice and barley), and is produced primarily in Yamanashi Prefecture, Japan. We characterized the microbiota of *Koshu miso* at three distinct fermentation stages. Our analysis revealed that the genus *Staphylococcus* dominated across all miso samples. Notably, *Staphylococcus* sequences in the miso matched those found in rice and barley *koji*, indicating the influence of raw ingredients on the initial microbial community. Additionally, analysis of the manufacturing environment suggested similarities between the environmental surfaces and miso, highlighting the importance of the manufacturing environment in serving as a medium for microbial transfer. These findings underscore the critical importance of both raw ingredients and manufacturing equipment in shaping the microbial composition and evolution of miso throughout the fermentation process.

## Introduction

Miso, a traditional Japanese fermented soybean paste, also known as an umami-rich condiment, is valued not only for its distinctive flavor, but also for its potential health benefits, such as probiotic properties and high antioxidant content (Cao *et al*., 2019). The production of miso involves a complex fermentation process, and the use of wooden barrels, though rare, is a highly valued traditional method. This unique approach enhances the fermentation environment, where the interplay of microorganisms plays a crucial role in developing its characteristic taste, aroma, and nutritional profile. Indeed, the microbial profile of miso has been reported to be different based on their ingredients (Kothe, Rasmussen, *et al*., 2024), as well as the fermentation environment, such as the temperature and the type of surfaces on which the microbial communities develop, in different breweries (Koide, Kanauchi and Hashimoto, 2024). This lies in contrast with other fermented foods, such as nukadoko, sake, and kimchi, that are dominated by Lactic Acid Bacteria at the end of the fermentation process (Ito *et al*., 2023; Niwa *et al*., 2023; Cha *et al*., 2024; Yamaguchi *et al*., 2024). The quality and uniqueness of fermented foods are closely linked to their microbial communities. Previous studies have shown that the microbial composition of fermented foods can be influenced by various factors, including raw materials, processing methods, and environmental conditions (Bokulich and Mills, 2013). The concept of “house microbiota” in food fermentation facilities has gained attention, suggesting that resident microbes in the manufacturing environment can significantly influence the microbial community and sensory properties of the final product (Bokulich, Lewis, *et al*., 2016; Imai *et al*., 2024). This concept is particularly relevant for traditional fermentation processes, where ancient practices may have selected for unique microbial ecosystems over time. In the case of miso, while some research has been conducted on its microbial composition (Harada and Yokoi, 2018), the specific influence of the manufacturing environment on the formation of the miso microbiome remains largely unexplored.

The study aims to investigate the microbial communities present on the raw materials and various equipment used throughout the stages of the miso fermentation process, and to assess how the microbial profiles of such manufacturing environments are related to the microbial community of the final miso product. We specifically analyzed *Koshu miso*, a unique type of miso made by mixing two types of *koji* (rice and barley) and produced primarily in Yamanashi prefecture, Japan. Assessing traditional practices and new manufacturing methods to maintain desired microbial profiles of miso has practical implications that allows us to gain insight into the microbial transfer from the manufacturing environment to the fermented foods. This research contributes to our understanding of miso fermentation, as well as to the broader field of microbiology in fermented foods.

## Materials and Methods

### Sample collection

Miso samples (Day 1, 10-month, 10-month miso in tube) were collected using individually wrapped, disposable plastic spoons and stored in 50 mL tubes. Samples of the manufacturing environment were collected from 10 locations, including the floor, raw materials, and equipment surfaces in the factory, by a swabbing method (Figure 1) between October 2023 to February 2024. Wooden barrel and stainless barrel samples were taken from the floor of the barrels. The description of the roles of each equipment is listed in Figure S1. All samples were collected at Gomi-shoyu (Yamanashi, Japan) between October and November 2023. Sterile cotton-tipped swabs (ESwab™; Copan Diagnostics, Brescia, Italy) were used for sampling, and swabs were stored in tubes with Liquid Amies Medium solution (Copan Diagnostics, Brescia, Italy). As for the raw ingredients, rice *koji* was prepared from *Shiro-Moyashi Tane Koji*, a fermentation starter obtained from Higuchi Matsunosuke Shoten (Osaka, Japan). Barley *koji* was prepared from a mixture of three fermentation starter cultures - *Mugi-miso Tane Koji*, *Shochu Ki-Koji*, and *Ogon* - from Higuchi Matsunosuke Shoten (Osaka, Japan), Akita Konno Co., LTD (Akita, Japan), and Hishiroku (Kyoto, Japan), respectively. All samples, including the miso, the manufacturing environment, and the raw ingredients, were immediately frozen and stored until DNA extraction.

**Figure 1.**
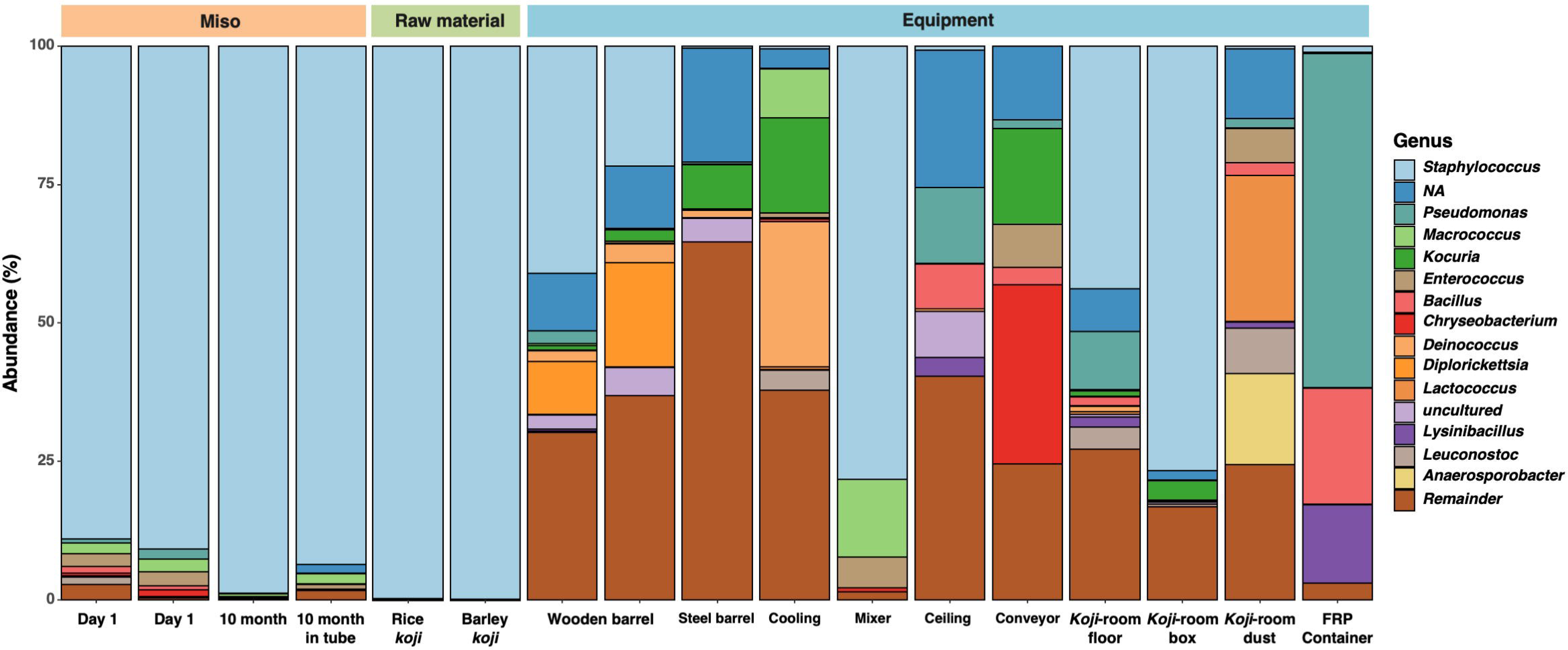
Genus-level taxonomic composition bar plots. The top 15 genera are listed, and the rest are noted as Remainder.

### DNA extraction and high-throughput sequencing

DNA isolation from 50 mL miso samples was performed using the bead-beating method using the multi-beads shocker (Yasui Kikai, Osaka, Japan). The extracted DNA was quantified using Qubit™ (Thermo Fisher Scientific, Massachusetts, USA). The DNA was further purified using a GenCheckⓇ DNA Extraction Kit Type S/F. The first PCR cycle was conducted using ExTaq HS DNA Polymerase, where specific primers 341F (5’-CCTACGGGNGGCWGCAG-3’) and 805R (5’-GACTACHVGGGTATCTAATCC-3’) of targeting the V3–V4 region of the 16S rRNA were amplified (Klindworth *et al*., 2013). Two μL of the first PCR product, purified using AMPure XP beads, served as a template for library preparation and the second PCR product was again purified using AMPure XP beads. 16S rRNA gene amplicon sequencing was performed using 460 bp paired-end sequencing on the Illumina MiSeq platform (Illumina Inc., San Diego, CA, USA) at FASMAC Co., Ltd. (Kanagawa, Japan).

### Microbiome analysis

Raw FASTQ data obtained from the Illumina MiSeq platform were imported into the QIIME2 platform (2022.11) as qza files (Bolyen *et al*., 2019). Denoising sequences and quality control were performed using the QIIME dada2, and finally, sequences were produced into amplicon sequence variants (ASVs) (Callahan *et al*., 2016). ASVs were assigned to the SILVA database’s SSU 138 using the QIIME feature-classifier classification scikit-learn package (Quast *et al*., 2013; Bokulich *et al*., 2018). The ASVs classified as mitochondria, chloroplast, or unassigned were excluded from the sequences. Subsampling was performed to reduce bias due to differences in sequence depth between samples. Weighted UniFrac distances were calculated, and the microbial community structure differences between the tree groups were visualized following a principal coordinate analysis (PCoA). Data were visualized using ggplot (version 3.4.0), ggsci (version 3.0.3), qiime2R (version 0.99.6) and ggprism (version 1.0.5).

## Results

### Microbial community profiles in miso and raw materials

The genus of *Staphylococcus* was dominant in the first day (89.01% and 90.82%), four-month (35.61%) and ten-month (98.74%) miso samples in wooden barrels. The genus of *Macrococcus* (43.55%) and *Enterococcus* (15.40%) were also frequently detected in four-month miso (Figure 1). Similarly, *Staphylococcus* was predominant in ten months of miso fermentation in 50 mL tubes (93.52%). In the miso of this study, fermentation was completed from the beginning of the fermentation stage to the end, with *Staphylococcus* remaining significant. The *koji* samples (rice and barley), one of the raw materials of miso along with soybeans, also displayed a high abundance of *Staphylococcus* (99.79% and 99.80%, respectively) genus.

### Microbial community profiles in the manufacturing environment

Analysis of environmental samples revealed distinct microbial communities across different manufacturing areas (Figure S1). The soybean conveyor, which comes into direct contact with steamed soybeans, has the most abundant genera including *Chryseobacterium* (32.37%), *Kocuria* (17.31%), and *Exiguobacterium* (11.92%). The cooler for steamed rice/barley has its unique mix, with *Deinococcus* (26.26%), *Kocuria* (17.25%), and *Dermacoccus* (9.52%) being the most abundant genera. The ceiling of the mixing room, where the rice/barley are covered with *koji*, exhibited *Pseudomonas* (13.59%), *Bacillus* (8.15%), and some genera from the *Frankeniaceae* family (8.90%). The mixer (to mix soybeans, rice malt, barley malt, and salt water), *koji* -room floor, and the *koji* -room box, all exhibited *Staphylococcus* (78.18%, 43.82%, and 76.69%, respectively) as the dominant genus. The wooden barrel is dominated by *Staphylococcus* (41.05% and 21.67%) and *Diplorickettsia* (9.56% and 18.84%). The stainless steel barrel is home to a different set of genera, with *Skermanella* (8.22%) and *Kocuria* (8.12%) being the most abundant. *Staphylococcus*, which formed a majority of the microbiota in the wooden barrel, only formed 0.36% of the total microbiota in the stainless steel barrel. Lastly, the FRP container, which is another facility for fermenting miso, is dominated by *Pseudomonas* (60.27%), *Bacillus* (20.99%), and *Lysinibacillus* (14.21%).

### Differences in microbial diversity between traditional and modern fermentation equipments

Principal Coordinate Analysis (PCoA) based on weighted UniFrac distances revealed a high degree of similarity among the microbiomes of raw materials, miso, and the mixer (Figure 2). Within the manufacturing environment, samples from the *koji* room (including boxes and floors) exhibited microbial profiles most similar to those of miso, followed by wooden barrels. Interestingly, while stainless steel barrels come into direct contact with miso during production, their microbial profile showed a level of dissimilarity to miso comparable to that of non-contact surfaces such as conveyors.

**Figure 2.**
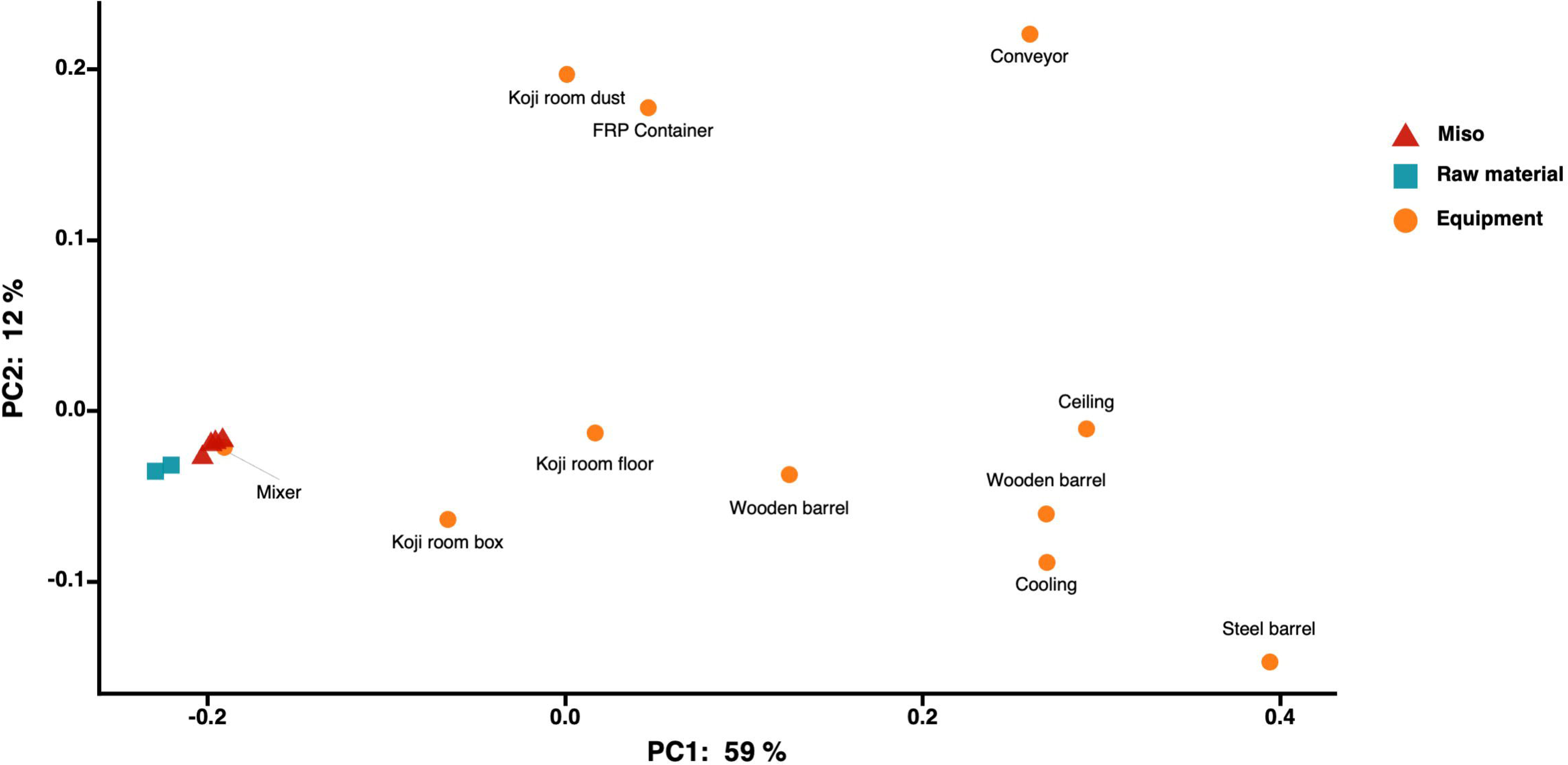
Principal-coordinate analysis (PCoA) by Weighted UniFrac distance of environmental samples and Miso samples. Data points represented by blue squares indicate the raw materials, orange circles represent the manufacturing environment, and red triangles represent the miso.

### Rice *koji* and barley *koji* were the main sources of the miso microbiome

We analyzed the distribution of Amplicon Sequence Variants (ASVs) for the three most abundant bacterial genera (*Staphylococcus*, *Macrococcus*, and *Enterococcus*) detected in four-month miso, focusing on their presence in raw materials and equipment (Figure 3). For *Staphylococcus*, eight ASVs were detected with 100 or more reads, with ASV1 predominating. Shared ASV analysis revealed that barley and rice *koji* contained the same ASV as the majority of Staphylococcus found in both the Day 1 miso and the final product. Notably, wooden barrel equipment also exhibited identical *Staphylococcus* ASVs. Within the manufacturing environment, the *koji* -room box and floor showed high abundances of ASV1 compared to other equipment. *Macrococcus* and *Enterococcus* ASVs were primarily associated with equipment in direct contact with raw soybeans, such as coolers and mixers (Table S1). *Macrococcus* ASV1 was most abundant in four-month miso (33,752 reads), followed by ten-month miso in tubes (2,088 reads). Other equipment, including wooden and steel barrels, showed smaller contributions, while rice and barley *koji* lacked *Macrococcus* ASVs entirely. Similarly, *Enterococcus* ASV1 was most prevalent in four-month miso (10,873 reads), with ten-month miso in tubes showing the second-highest abundance (1,015 reads). *Enterococcus* was detected on various equipment, including wooden barrels and conveyors. *Koji* -room dust exhibited a diverse range of Enterococcus ASVs, spanning seven distinct variants.

**Figure 3.**
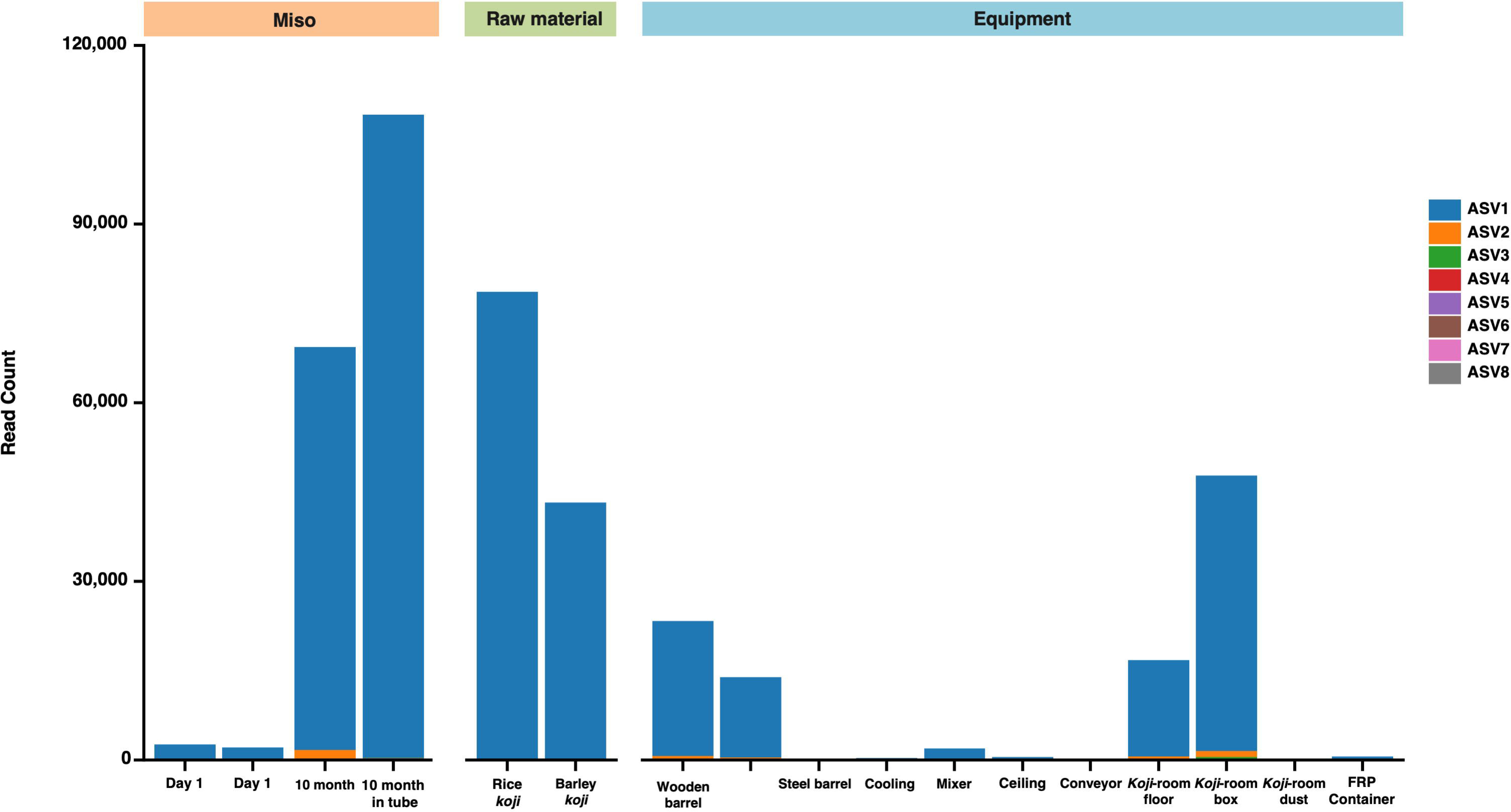
Stacked bar plot of amplicon sequence variants (ASVs) of *Staphylococcus*.

## Discussion

The consistent high abundance of *Staphylococcus* in both *koji* and miso samples throughout various fermentation stages underscores its potential significance in the miso fermentation process. Notably, ten-month miso samples exhibited a high abundance of *Staphylococcus*, regardless of whether they were fermented in wooden barrels or stored in plastic tubes after the first day. This persistent presence of *Staphylococcus*, even in non-fermentation vessels, suggests its crucial role in maintaining miso stability and quality over extended periods. These findings align with previous studies that have identified *Staphylococcus* as a key player in various fermented foods (Coton *et al*., 2010). *Staphylococcus* species are known to secrete enzymes capable of breaking down complex sugars and starches (Lakshmi *et al*., 2013), potentially enabling them to outcompete other microorganisms and dominate the microbiota in the final miso product (Kothe, Carøe, *et al*., 2024). Furthermore, the presence of *Staphylococcus saproph yticus*, renowned for its proteolytic and lipolytic activities, may contribute to the development of flavor compounds during fermentation (Kothe, Rasmussen, *et al*., 2024). Our observations are consistent with a previous study involving miso made with rice *koji*, soybeans, and yellow peas, which similarly reported a high abundance of *Staphylococcus*. This resemblance to our experimental miso, also made with soybeans and rice *koji*, further supports the potential universal role of Staphylococcus in miso fermentation across various recipes and production methods. While miso is known to contain high salt concentrations, usually within the range of 10-12%, certain *Staphylococcus* species like *Staphylococcus equorum* exhibit halotolerance, allowing them to thrive under saline conditions (Vega *et al*., 2019). It is important to note, however, that some previous studies on *koji* have reported different microbial compositions. For instance, a previous study did not detect *Staphylococcus*, instead identifying species such as *Bacillus*, *Weizmannia*, *Ochrobactrum*, and *Tetragenococcus* (Allwood, Wakeling and Bean, 2023). In another study (Hui *et al*., 2017), *Ochrobactrum* species accounted for over 99% of the total bacterial sequences across three different *koji* samples. These discrepancies may be attributed to regional variations in manufacturing practices and differences in raw ingredients (rice and barley *koji*), underscoring the need for comprehensive comparative studies across diverse manufacturing environments.

Our Principal Coordinate Analysis (PCoA) results indicate similar microbial compositions between the first-day miso samples and the raw ingredients, suggesting the foundational role of *koji* in shaping the initial miso microbiome (Figure 2). The shared fermentation starter used in both rice and barley *koji* likely contributed to the initial microbial composition of day 1 miso. Considering that both rice and barley are washed, steamed, and inoculated with the *tane-koji* fermentation starter, the microbial communities of these *koji* most likely derive from *tane-koji*.

Interestingly, the mixer also exhibited a microbial profile similar to the raw ingredients, possibly due to its direct contact with rice and barley malt during the fermentation process. Furthermore, the *koji* -room box and floor displayed microbial profiles relatively similar to the raw ingredients. This observation suggests that the manufacturing environment may serve as a reservoir for microbial transfer to miso, highlighting the potential importance of environmental factors in shaping miso microbiomes throughout the fermentation process.

In addition, the wooden barrel floor exhibits a more similar profile with miso, compared to other fermentation vessels such as the stainless steel surfaces. While we have previously identified the abundance of *Staphylococcus* as a key characteristic of miso fermentation in this study, *Staphylococcus* was detected at significantly low abundance in the steel barrel surfaces (0.36%) compared to the wooden barrel surfaces (41.05% and 21.67%) (Figure 1). The significantly higher abundance of *Staphylococcus* on the wooden barrel surfaces may suggest that the wooden barrel environment may help maintain a more natural and diverse microbiome by preserving and enhancing the microbial composition, initially introduced by the raw ingredients. The importance of utilizing traditional fermentation vessels is highlighted by a previous study conducted on Lambic beer. Here, the barrels used during the manufacturing process exhibited some yeast species found in the final beer product, suggesting the role of the wooden barrels as a microbial inoculation source (Bongaerts, De Roos and De Vuyst, 2021). In an additional research on the effect of sanitization of the wooden barrels on the abundance of microorganisms, microorganisms detected on the interior surfaces of the wooden barrels mostly reflected the microbiota present in the Lambic beer they contained previously (De Roos, Van der Veken and De Vuyst, 2019), suggesting a complex interplay of the exchange and growth of microorganisms between the wooden barrels and the fermented product. This difference may be due to the porous nature of wood, which can harbor a more diverse microbial community (Cappello *et al*., 2017). In this study, however, the microbial compositions of 10-month miso fermented in wooden barrels and 10-month miso stored in tubes were relatively similar, dominated by the same *Staphylococcus* ASVs. These results highlight the importance of further research on the impact of the wooden barrel manufacturing environment.

To identify the origins of the *Staphylococcus* genus identified in the final miso product, we analyzed the ASVs of the *Staphylococcus* genus and found that the sequences identified in rice and barley *koji* samples matched the sequences of the *Staphylococcus* genus in the miso samples (Figure 3). While the miso exhibits a more diverse microbial profile at four months into the fermentation stage, *Staphylococcus* from the raw ingredients dominates the microbiota by the end of the ten-month period. Indeed, the wooden barrel and both samples of the ten-month miso exhibited the same *Staphylococcus* ASV1 found in rice and barley *koji*, suggesting that the unique microbial composition of miso is preserved throughout the fermentation process.

In addition, the *koji* -room box and the *koji* -room floor showed high abundances of *Staphylococcus*, further suggesting that the interplay between the raw ingredients (rice, barley, and *tane-koji)* and the manufacturing environments serve a role in microbial transfer to miso. While the *Staphylococcus* genus formed a majority in the microbiota, other genera including *Macrococcus* and *Enterococcus* were commonly observed in all miso samples, notably the four-month miso. While the reason for a temporal increase in microbial diversity remains unclear, the cooler and the mixer showed a high presence of *Enterococcus* and *Macrococcus*. The higher abundance of *Enterococcus* in four-month miso is consistent with previous studies identifying *Enterococcus* sp. in traditional fermented foods (Onda *et al*., 2003; Fugaban *et al*., 2021). Another study has also reported the presence of *Enterococcus faecium* during the soaking process of soybeans (Elhalis, Chin and Chow, 2024), which explains the presence of *Enterococcus* ASV1 in the soybean conveyor, the cooler and the mixer, which are all equipment that come into direct contact with soybeans, one of the raw ingredients of miso. Indeed, despite their absence in the raw materials such as rice and barley *koji*, *Enterococcus* may be introduced during the early stages of miso production, where they are nurtured and eventually contribute to the fermentation process.

Similarly, while *Macrococcus* may not derive from soybeans or the raw ingredients, as it was not detected in the conveyor and rice and barley *koji*, it is possible that *Macrococcus* is introduced during the handling of soybeans in the cooler and mixer, where conditions may favor its growth. This suggests that while *Macrococcus* is not present in the raw ingredients, it could be introduced during the early stages of miso production, particularly through equipment that comes into direct contact with the soybeans. This introduction of microorganisms from the manufacturing environment could have a profound impact on the quality and characteristics of the final miso product. For example, *Macrococcus caseolyticus*, the most commonly studied species of *Macrococcus*, is often associated with the development of aroma and flavor in fermented foods (Ramos, Vigoder and Nascimento, 2021). Understanding the roles of these microbes is essential for controlling the consistency and quality of miso, as they could influence its texture, taste, and even potential health benefits. Thus, the persistence of the *Macrococcus* and *Enterococcus* genera further highlights the role of not only the raw ingredients in shaping the microbiome of miso, but also the importance of the transfer of bacterial strains between fermented foods and specific manufacturing environments.

Such transfers of environmental microbes to the final miso product highlights the concept of “terroir” in fermented foods, where the local environment imparts the unique characteristics of the product (Bokulich, Collins, *et al*., 2016). In addition, the interactions between humans and the microbial community of the built-environment (MoBE) is known to create a unique microbial ecosystem (光平, 2023). These points suggest that the traditional manufacturing environment contributes significantly to the distinctive qualities of artisanal miso.

## Conclusion

In conclusion, our results demonstrate that the formation of a unique microbial community in the final miso product begins with the characterization of the initial miso microbiota dominated by *Staphylococcus*. We have also identified the sources of *Staphylococcus* as the raw ingredients, rice and barley *koji*, which used a common fermentation starter. In addition, the results of the ASVs analysis of *Staphylococcus*, *Macrococcus*, and *Enterococcus*, suggest that equipment from the manufacturing environment, such as the cooler and the mixer, also shape the microorganisms of miso by serving as major reservoirs for microbial transfer to miso. This study provides a scientific basis for the value of traditional manufacturing methods and opens new avenues for understanding and potentially improving the quality of not only miso, but all kinds of fermented foods.

## Limitations

The analysis provides a snapshot of the microbial community and does not capture temporal dynamics throughout the fermentation process. The study is constrained by a small sample size and lacks statistical testing, making the suggested microbial transfer between *koji* and miso only a tentative explanation for an observation. Future research should incorporate time-series sampling to elucidate the succession of microbial populations during fermentation. With regards to shared ASV analysis, it must be noted that shared ASVs are limited by the shortcomings of 16S rRNA amplicon sequencing. To analyze which specific bacteria have been transferred, it is necessary to conduct whole-genome analysis and use metagenomics to compare the results in a more comprehensive manner.

## Data availability

The datasets generated through 16S rRNA amplicon sequencing are available and deposited in the DDBJ Sequence Read Archive (SRA) database under accession numbers DRR597132-DRR597149 and BioProject PRJDB18832.

## Acknowledgements

All authors thank Morgenrot Inc. for providing the computational environment for the analysis and Gomi-shoyu for collecting samples and giving advices for study design.

## Ethics declarations

### Author Contribution

KI designed the experiments, collected samples, performed microbiome analysis. KI and MY drafted the original manuscript. All authors have contributed to the manuscript and approved the submitted version.

---

**Figure S1**. An overview of multiple processes for miso fermentation with sampling targets in the study.

**Table S1**. The number of amplicon sequence variants (ASVs) of *Macrococcus* and *Enterococcus*.

