## Supplementary figures and images for "Raw materials and manufacturing environment as determinants of miso microbial community"

### Figure S1

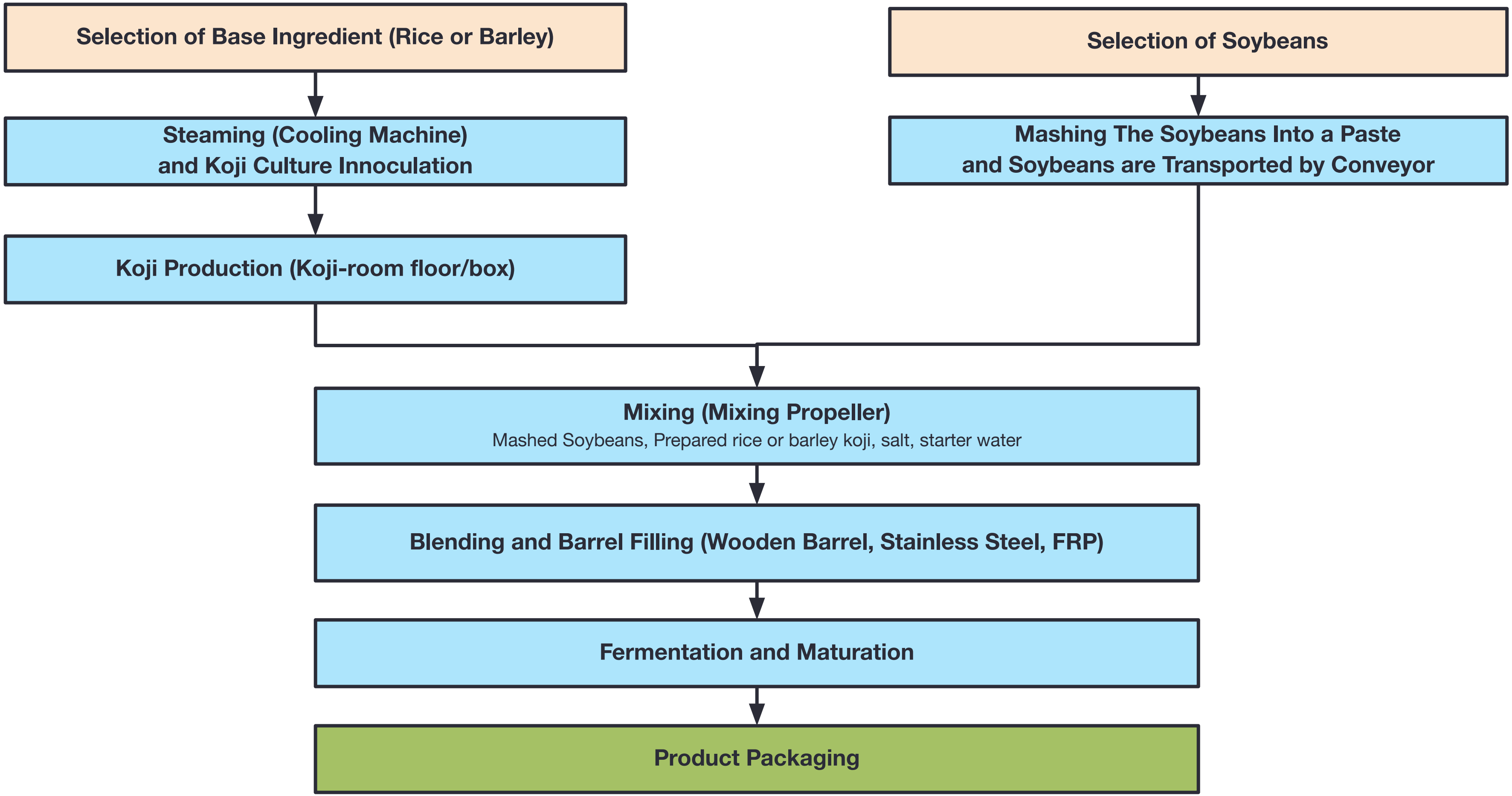

### Figure S2

*Macrococcus*

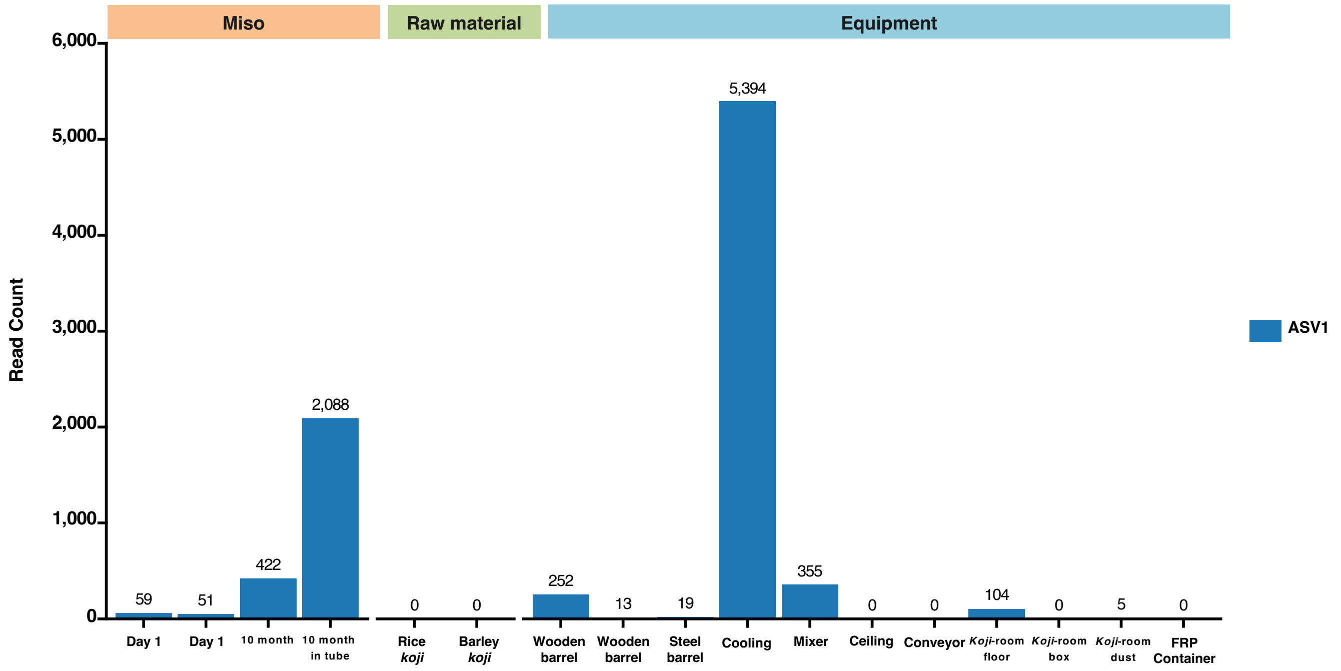


*Enterococcus*

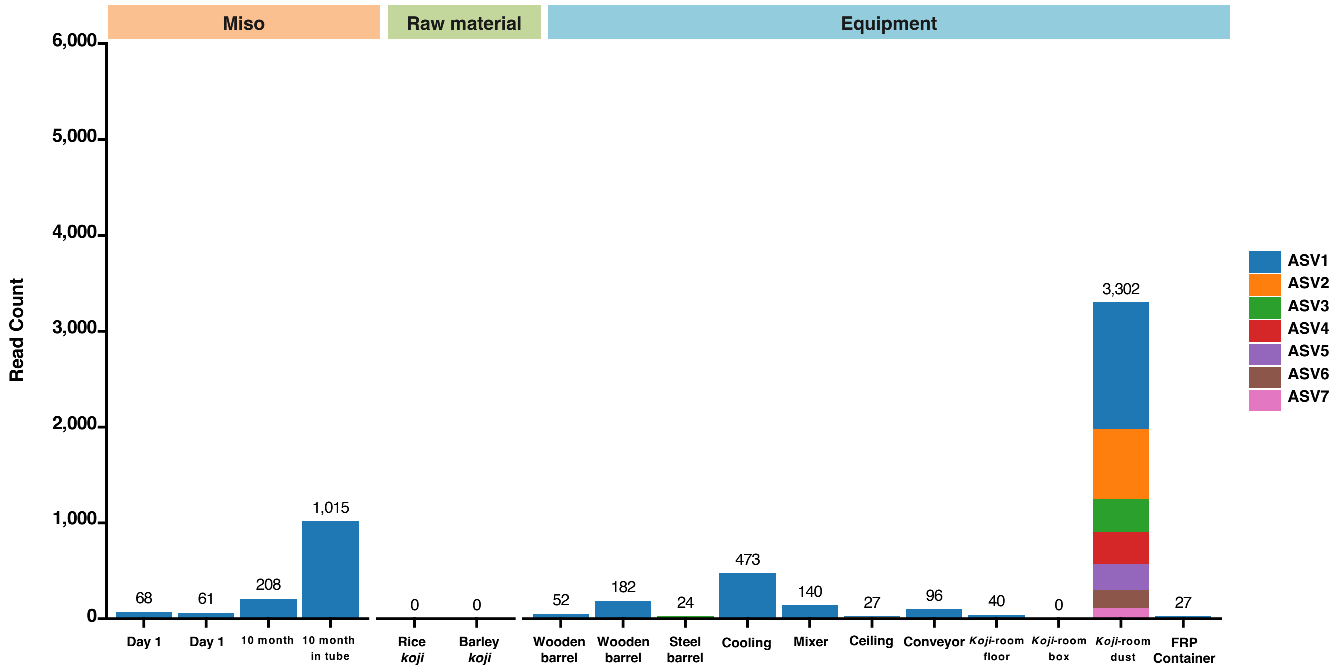
